# Histological techniques for the visualisation and identification of polymicrobial communities in mucosal tissue

**DOI:** 10.1101/2025.11.26.690509

**Authors:** Harriet J. Giddings, Ana Teodosio, Jordanne Jones, William Butterworth, Claire D. Shannon-Lowe, Jeffrey A. Cole, Amanda E. Rossiter-Pearson

**Affiliations:** Department of Microbes, Infection and Microbiomes, School of Infection, Inflammation and Immunology, College of Medicine and Health, University of Birmingham, UK; Institute of Microbiology and Infection, University of Birmingham, UK; Birmingham Tissue Analytics, University of Birmingham, UK; Department of Immunology and Immunotherapy, School of Infection, Inflammation and Immunology, College of Medicine and Health, University of Birmingham, UK; School of Biosciences, College of Life and Environmental Sciences, University of Birmingham, UK

**Keywords:** Helicobacter pylori, histology, gastric adenocarcinoma, microbiome, spatial biology

## Abstract

Sequencing studies have generated masses of data linking mucosal bacteria to health and disease, particularly in organs such as the colon, mouth and stomach. However, extracting microbial DNA from mucosal sites with lower bacterial biomass can introduce significant variation given the heterogeneity of bacterial colonisation within pathological microenvironments. Therefore, a histological technique that identifies mucosal samples with high microbial mass samples prior to applying sequencing technologies, would increase reproducibility and reduce issues of low signal: noise ratios commonly observed during 16S rRNA gene sequencing. Using RNAscope technology, we have previously shown that non-*H. pylori* bacteria invade the gastric lamina propria in *H. pylori-*infected patients diagnosed with chronic gastritis and gastric intestinal metaplasia. However, RNAscope technology is costly and time consuming. Here, using gastric tissue from patients with *Helicobacter pylori*-associated precancer and cancer, we applied the use of a modified Gram stain technique to identify gastric tissue with high bacterial biomass. We validated the presence of bacteria using custom-designed RNAscope probes on consecutive tissue sections and then we visualised bacteria within a polymicrobial community at the Phylum, Kingdom and Genus taxonomic levels. This study validates the cost-effective modified Gram stain as an efficient way to select samples with high microbial biomass for downstream applications investigating host-microbiota relationships.

## Introduction

*Helicobacter pylori* has been designated a class one carcinogen because it is associated with 70% of gastric adenocarcinoma (GAC) cases (1). GAC follows a progressive cascade that initiates with chronic gastritis, proceeds via the precancerous stage of gastric intestinal metaplasia (GIM), dysplasia and finally GAC (2). Despite the strong association between *H. pylori* and GAC, many questions remain unanswered. Multiple sequencing studies have shown that during progression to gastric cancer, *H. pylori* load decreases and appears to be replaced by other bacteria (3, 4). However, the mechanistic link between these observations is, at best, only partially resolved. Given that sequencing studies lack spatial resolution, we applied spatial biology technologies to resolve the gastric microbial community structure in chronic gastritis and precancer tissue sections (5). We found that non-*H. pylori* bacteria invade the gastric lamina propria in *H. pylori-*associated chronic gastritis and GIM. We also showed the preliminary application of a modified Gram stain to identify these invasive bacteria in histological gastric GIM tissue sections.

Currently, Haematoxylin and Eosin (H&E) staining is routinely used for the combined clinical diagnosis of *H. pylori* infection and associated pathology, such as GIM. However, the H&E stain is unsuitable for revealing bacterial complexity in mammalian tissues because Gram-positive bacteria are not visible against the staining properties of mammalian cells. The Gram stain is exceptionally useful for the preliminary characterisation of bacteria in many types of sample.

There is much controversy surrounding the presence of intratumoral microbiomes, particularly within internal organs not of the gastrointestinal tract. Indeed, a report indicating distinct microbial signatures within tumours from a range of organs (6, 7) was later retracted due to fundamental flaws in incorrectly annotating human sequences as bacterial (8). In addition to the biological consequences of incorrectly assigning microbiomes to tumour types, these studies highlight the necessity to use complementary imaging or culture techniques to validate sequencing data. Nonetheless, the question of intratumoral microbiomes and their role in carcinogenesis and response to treatments is an intense and ongoing field of research. Indeed, intracellular bacteria have been identified in a range of tumour types (9). With the advent of high-resolution spatial biology approaches, it is now imperative that more accessible methods are available to select the most appropriate biopsy samples for more detailed and comprehensive analyses, using complementary culturing, sequencing and imaging technologies to resolve the strain-level identity of colonising microbes.

Here, we compare the use of H&E, modified Gram stain and RNAscope *in situ* hybridisation histological tools for the visualisation of *H. pylori* and non-*H. pylori* bacteria on gastric chronic gastritis, GIM and cancer tissue sections. We now show that with only minor modifications, the Brown and Hopps (1973) method can reveal wide variations in the localisation of bacteria in gastric tissue samples from different patients. The preliminary microscopic examination has enabled us to select samples suitable for targeted genus-specific staining. This report evidences the use of modified Gram staining for the identification of samples suitable for further studies, which can be applied to research surrounding the role of the microbiome in carcinogenesis and within intratumoral tissue.

## Materials and methods

### Preparation of FFPE tissue sections for staining

Archived gastric tissue samples were collected from consenting patients at the Queen Elizabeth Hospital, Birmingham (Ethics #17–285) and prepared at the Human Biomaterial Resource Centre (HBRC), University of Birmingham, by fixing formalin and embedding in paraffin (FFPE). 4 μm thick tissue sections were then kindly prepared for staining by the HBRC or by Dr Gary Reynolds (Department of Cancer and Genomic Science, University of Birmingham). Prior to all staining methods, each section was baked for 1 hour at 60°C before incubating three times in xylene for 5 minutes, followed by a series of 3-minute incubations in ethanol at 100, 95, 75 and 50% concentration.

### Conventional Gram stain of FFPE tissue sections

Dewaxed tissue sections were incubated in crystal violet solution for 2 minutes, rinsed with distilled water and immediately incubated in Gram’s iodine for 5 minutes. Sections were then further rinsed and immediately decolourised with acetone for 10 seconds, before dipping in distilled water. Gram’s safranin was applied to the tissue sections for a further 5 minutes, before sections were rinsed twice more in distilled water, incubated in xylene for 3 minutes and immediately mounted in DPX Mountant with a glass cover slip. Sections were air dried for 48 hours and viewed using a light microscope. Whole slide scans were also obtained using a Zeiss Aperio Whole Slide Scanner.

### Modified Gram stain of FFPE tissue sections

Consecutive, dewaxed tissue sections were incubated in crystal violet solution for 2 minutes, rinsed and incubated in Gram’s iodine for 5 minutes. Sections were rinsed and immediately decolourised with acetone for 10 seconds, before dipping in distilled water immediately followed by incubation in Gram’s safranin for a further 5 minutes. Following addition of all stains, the sections were rinsed twice more and incubated in Gallego solution (distilled water, 0.05% formaldehyde, 0.01% acetic acid) for a further 5 minutes. Sections were then rinsed, and sequentially dipped 5 times in acetone, 0.05% picric-acid acetone, acetone-xylene and xylene in a fume hood. Finally, sections were blotted dry and mounted in onto a glass microscope slide in DPX Mountant with a cover slip. Sections were air dried for 48 hours and viewed using a light microscope. Whole slide scans were also obtained using a Zeiss Aperio Whole Slide Scanner.

### Manual FFPE tissue preparation

Consecutive, dewaxed sections were briefly washed in distilled water before incubating in RNAscope hydrogen peroxide for 10 minutes at room temperature. Sections were rinsed in distilled water and placed in heat-proof Simport staining jars containing RNAscope antigen retrieval buffer, whereby heat-induced antigen retrieval (HIER) was conducted with a microwave oven at 20% power for 15 minutes. Sections were then briefly added to 100% ethanol and dried at 60°C for 5 minutes, prior to addition of RNAScope protease III for 30 minutes at 40°C in a humidified box.

### FFPE RNAscope in situ hybridisation and immunohistochemistry

Methods described in Giddings *et al*. were followed for combined RNAscope *in situ* hybridisation and IHC. Briefly, sections were washed twice in RNAscope wash buffer, prior to following the RNAscope multiplex Fluorescent v2 Assay according to manufacturer’s instructions. Sections were incubated with the indicated RNAscope probes shown in Table 1 for 2 hours. Sections were washed twice more in RNAscope wash buffer and incubated overnight in 5x sodium saline citrate (SSC) buffer at room temperature. On day two, sections were removed from SSC buffer and washed twice in RNAscope wash buffer, prior to continuing the v2 assay involving amplification and labelling of all RNAscope probes. Immediately following the v2 assay, E-cadherin and Mucin 2 was fluorescently labelled using primary antibodies (Table 1) and the Opal 7-colour IHC detection kit, counter-stained with DAPI and mounted. Sections were then visualised using a Mantra 2 Quantitative Pathology Workstation at 4x magnification and whole slide scans were obtained with a Akoya PhenoImager HT whole-slide scanner.

**Table 1.**
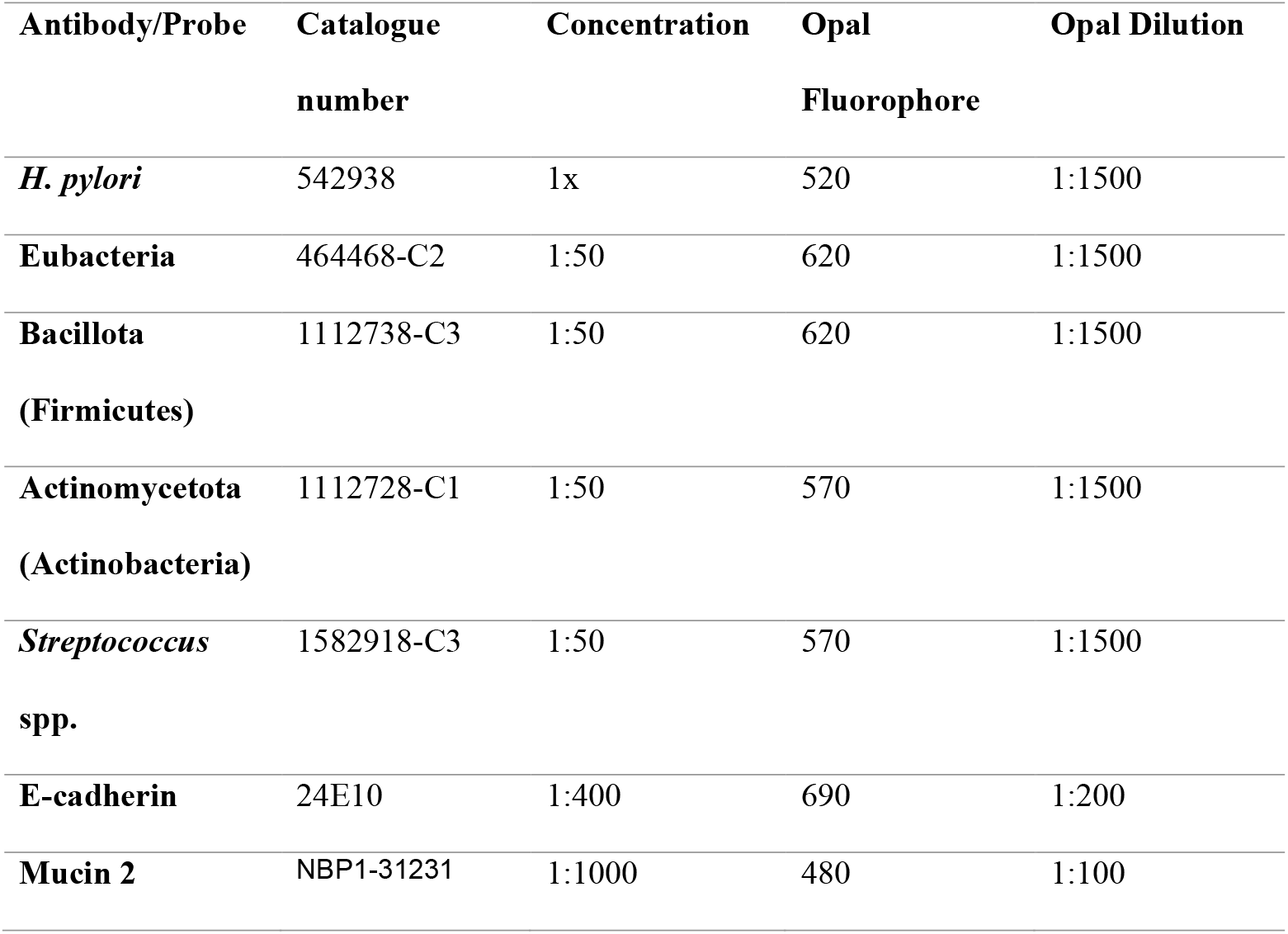
Antibodies and probes used for automated RNAscope IHC assays.

### Custom-design of RNAscope probes

RNAscope probe design algorithm examines each oligo probe in the probe set for potential cross-reactivity against the host transcriptome to ensure the probe specificity. There is a verification procedure following each major step during probe design to guarantee accuracy. First, the probe design algorithm selects oligos complementary to the input target sequence with compatible melting temperature for optimal hybridisation and minimal cross-hybridisation to off-target sequences. Next, during the synthesis and formulation process, oligo synthesis QC is performed on each oligo probe before providing design feasibility. Further design information is available on the Bio-techne website (http://www.bio-techne.com).

## Results

### Comparison of H&E, modified Gram stain and RNAscope *in situ* hybridisation for the detection of bacteria in gastric tissue sections

Given that H&E staining is currently used for the detection of *H. pylori*, we compared H&E staining with modified Gram staining for the visualisation of bacteria in gastric tissue. To validate bacterial localisation, we also used RNAscope *in situ* hybridisation (ISH) for the detection of *H. pylori* and non-*H. pylori* bacteria, using probes against the conserved or variable regions of the 16S ribosomal RNA gene, respectively (5). Sections of gastric punch biopsy tissue from a patient with *H. pylori-*positive chronic gastritis (Figure 1A-C) or gastric adenocarcinoma (Figure 1 D-F) were stained using either the traditional Gram stain or the modified stain. Figure 1A shows the detection of *H. pylori* within a gastric gland, whereas *H. pylori* could not be visualised using the modified Gram stain on a consecutive section. Figure 1C shows the clear visualisation of *H. pylori* in the same gland using RNAscope ISH. In contrast, Figure 1E shows that modified Gram staining is superior to H&E staining (Figure 1D) for the detection of non-*H. pylori* bacteria given that a variety of Gram-positive and negative bacteria were detected within this microenvironment. Again, RNAscope ISH confirmed the presence of non-*H. pylori* bacteria within this region (Figure 1F).

**Figure 1.**
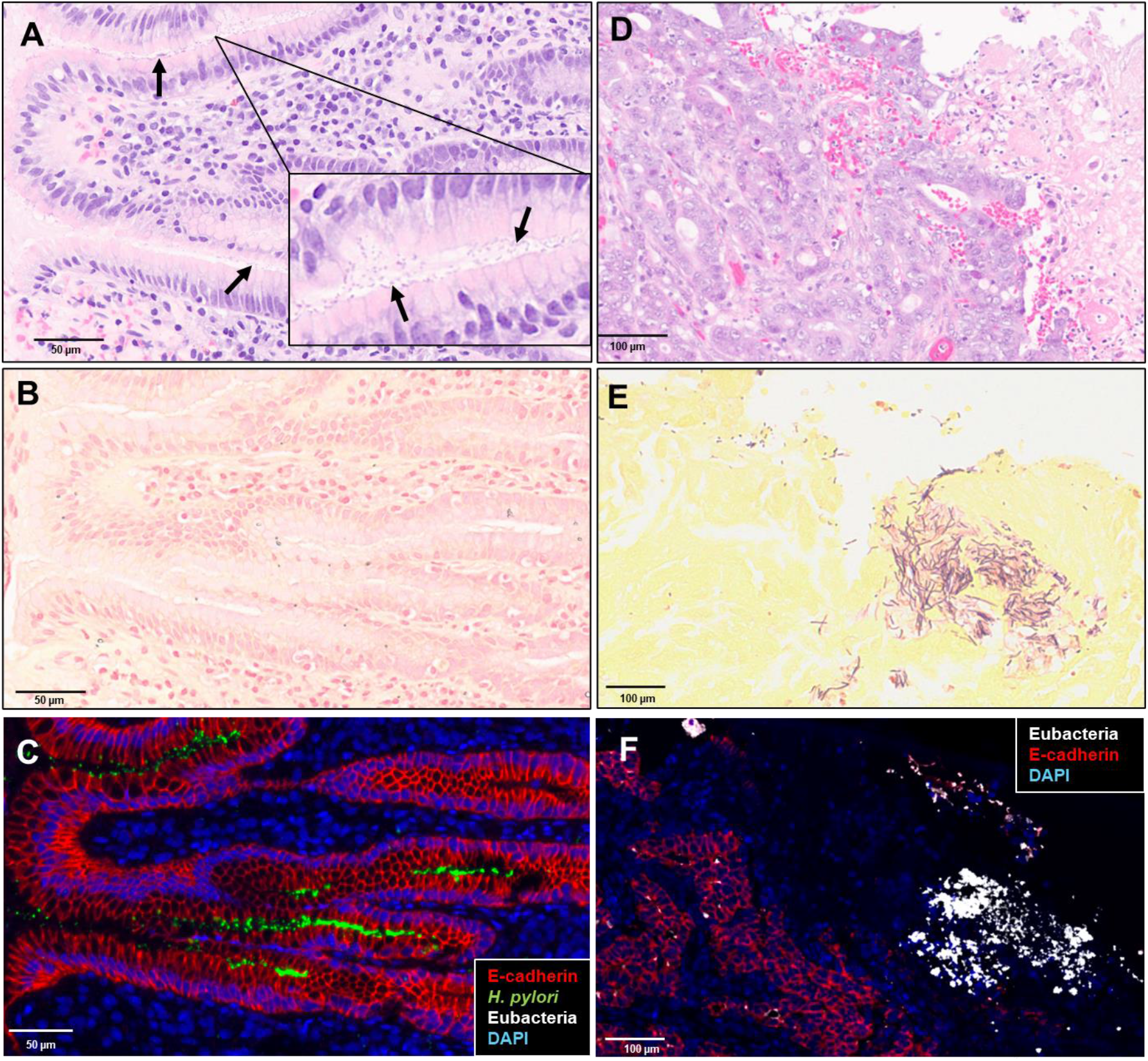
Comparison of H&E, modified Gram stain and RNAscope *in situ* hybridisation for the detection of *H. pylori* and the gastric microbiome in gastric tissue sections. FFPE tissue sections from a patient with *H. pylori*-positive chronic gastritis (A-C) or gastric adenocarcinoma (D-F) were deparaffinised and rehydrated. Sections were stained with H&E (A&D) or modified Gram staining (B&E) and visualised at 1000x magnification by light microscopy with oil immersion. Consecutive sections were also labelled with an RNAScope probe against specific or conserved regions of the 16S rRNA gene to detect *H. pylori* (green) or Eubacteria (white), respectively (C&F). Immunofluorescence was also used to detect E-cadherin (red). Sections were counterstained with DAPI and a Akoya PhenoImager HT whole slide scanner was used for visualisation at 40x magnification.

### Bacterial diversity revealed by modified Gram stain and RNAscope probes in gastric tissue sections

The initial microscopic examination of tissue samples from the GAC patient (Figure 1D-F) merited more detailed analysis of the diversity, abundance and distribution of bacteria in gastric tissue. Additionally, given that Figure 1 used gastric tissue from patients at the start and the end of Correa’s carcinogenic cascade, we also sought to identify to apply this technique to a patient with GIM, the middle stage of Correa’s cascade. GIM is a pre-cancerous condition of the stomach in which stem cells aberrantly differentiate to intestinal goblet cells that secrete the intestinal-specific Mucin 2. We identified a large region of bacterial colonisation in this GIM patient tissue section, using the Eubacteria probe (Figure 2A), which was also surrounded by a layer of Mucin 2. The diversity of this community was confirmed using custom-designed RNAscope probes that target bacteria belonging to the Bacillota (formerly named Firmicutes) or Actinomycetota (formerly named Actinobacteria) Phyla. Staining with these probes showed a dominance of Firmicutes and a smaller load of bacteria belonging to the Actinomycetota phylum (Figure 2B). The unstained regions within the community shown in Figure 2A likely represent Bacteroidota (formerly named Bacteroidetes) given their staining with the Eubacteria probe, but not with either the Bacillota or Actinomycetota probes. The H&E staining of the consecutive tissue section confirmed a high bacterial load, as revealed by purple and pink-stained regions within the same region (Figure 2C). To further confirm the use of the modified Gram stain on cancer tissue sections to reveal cellular diversity, we used a modified Gram staining of the GAC patient tissue shown in Figure 1D-F. Regions of bacterial colonisation within the tissue were confirmed with RNAscope ISH staining against Eubacteria (Figure 3A). Bacterial diversity was revealed by modified Gram staining of a consecutive tissue section and can be seen in Figure 3B-E. These reveal a range of Gram-positive cocci, Gram-negative spindle-shaped bacteria and Gram-negative rods. Given the recent interest in the role of *Streptococci* spp. in the development of GAC (10), we also stained consecutive tissue sections with the Bacillota probe of which *Streptococci* belong to. A significant proportion of the bacteria detected with the Eubacteria probe, shown in Figure 4A, stained with the Bacillota probe (Figure 4B). However, a large region of bacteria did not stain with the Bacillota probe, indicating further diversity at the Phyla level. To further interrogate the abundance of *Streptococcus* spp., an additional custom-designed genus-level probe targeting, multiple *Streptococci* species was used to stain a consecutive tissue section. As can be seen in Figure 4C, a relatively small region of the Bacillota population was stained with the *Streptococcus* probe, indicating that there are multiple other bacteria within this phylum that colonise the GAC tissue, rather than solely *Streptococci*.

**Figure 2.**
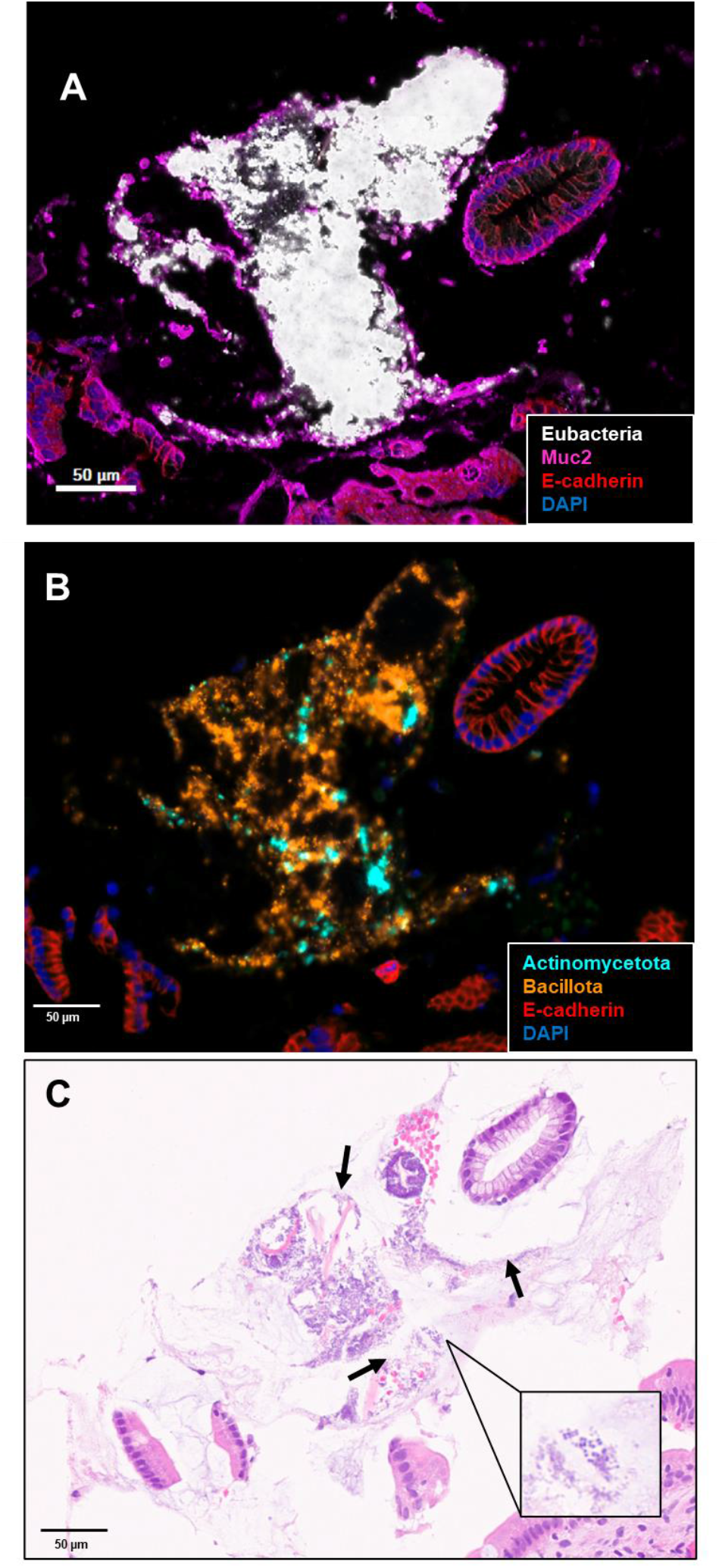
Phyla-specific RNAscope compared with H&E for detection of high bacterial biomass in gastric tissue sections. Punch gastric biopsy FFPE sections from an archived *H. pylori*-negative GIM patient were obtained from the HBRC, deparaffinised and rehydrated. (A) Sections were labelled with E-cadherin (red), an RNAscope probe against Eubacteria (white) (A) or Actinomycetota (cyan) and Bacillota (orange) (B). Both sections were counterstained with DAPI and a Akoya PhenoImager HT whole slide scanner was used for visualisation at 40x magnification. (C) H&E was used for staining and the section was visualised at 1000x magnification by light microscopy with oil immersion.

**Figure 3.**
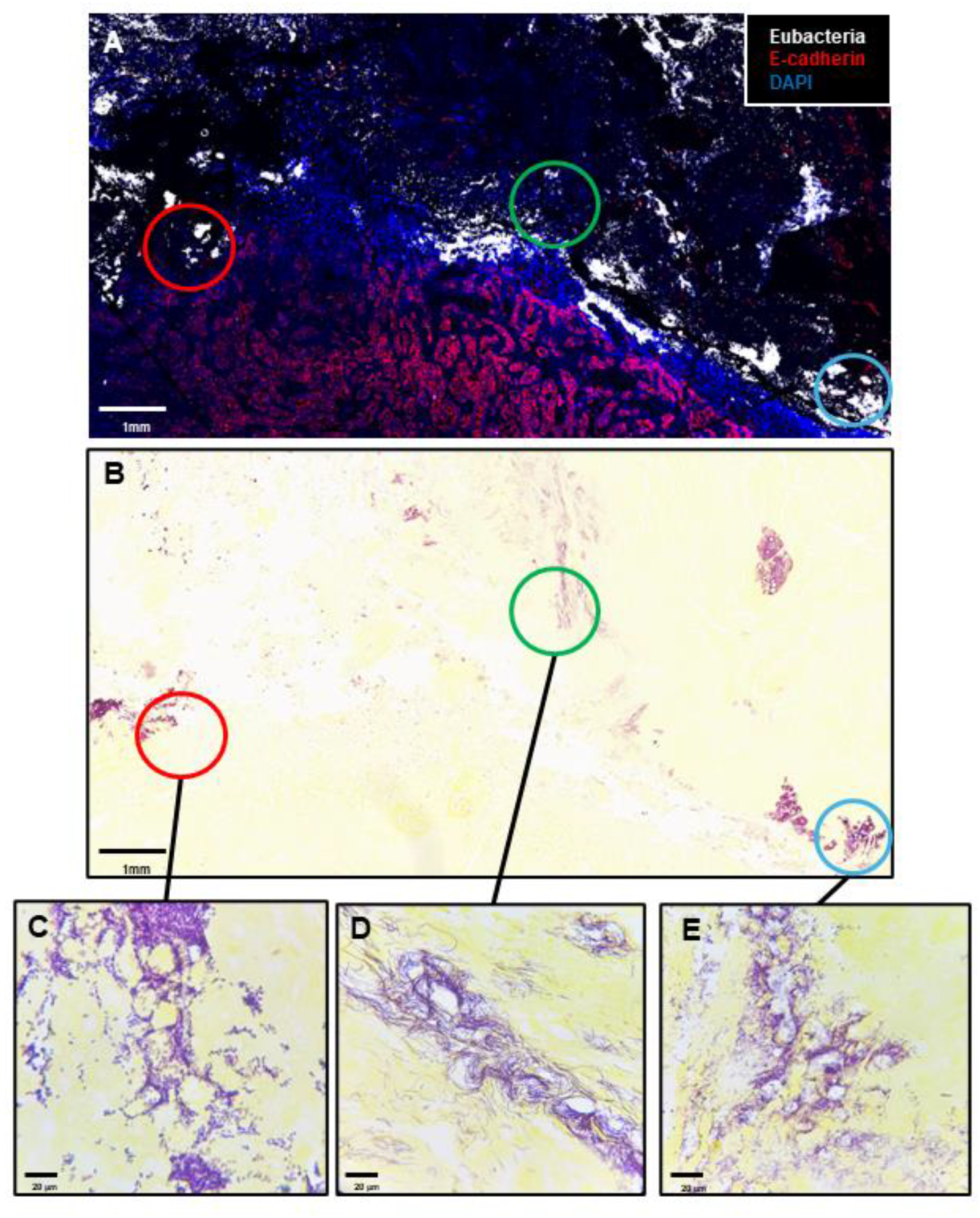
Modified Gram stain of gastric adenocarcinoma tissue sections. Consecutive sections of an archived FFPE gastric adenocarcinoma patient were prepared for staining by dewaxing in xylene, rehydrating in ethanol and rinsing in distilled water. (A) RNAscope probes against Eubacteria (white) and an antibody against E-cadherin were used for staining followed by counterstaining with DAPI and visualisation with a Akoya PhenoImager HT whole slide scanner at 40x magnification. (B) Modified Gram staining was visualised at 4x using a light microscope and (C-E) show microniches containing different bacterial morphologies visualised at 1000x magnification under oil immersion.

**Figure 4.**
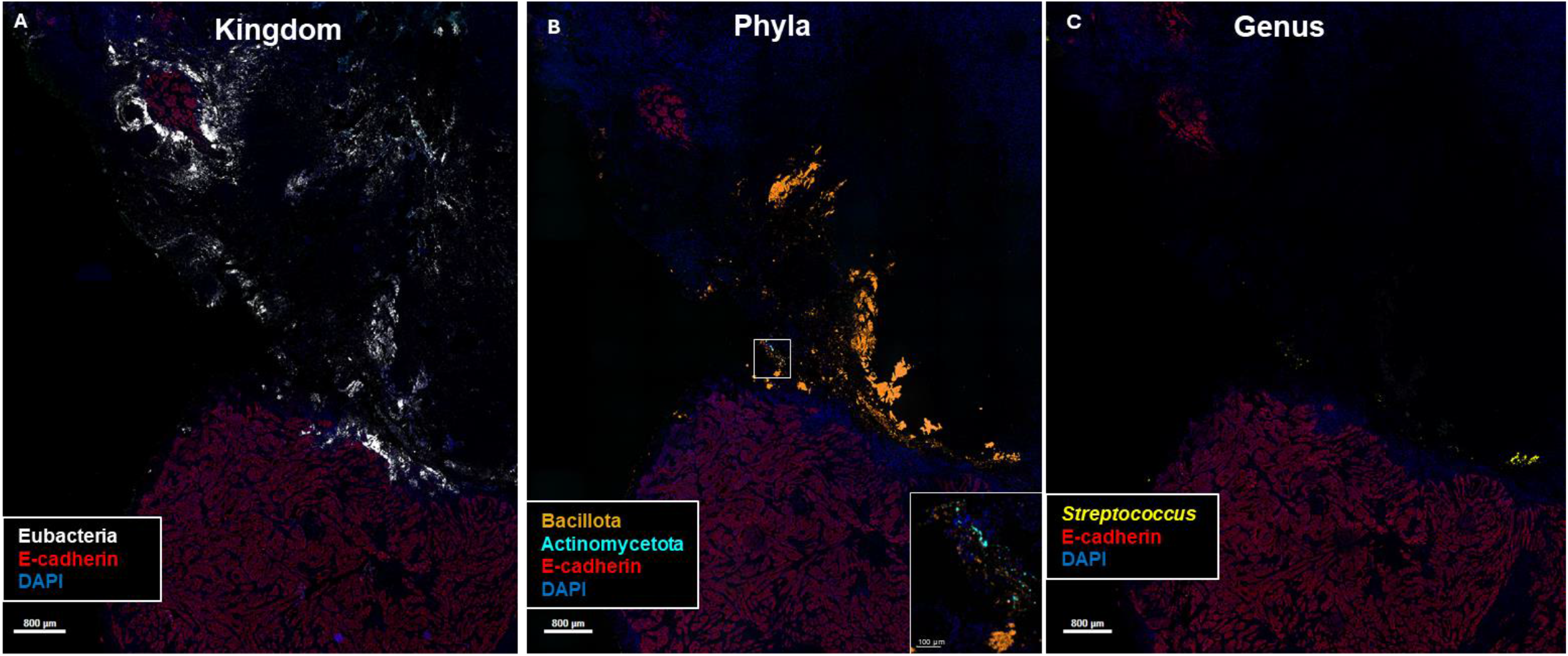
Phyla and genus-specific probes can identify bacterial colonisation of GAC tissue by *Streptococcus* spp. Consecutive FFPE GAC tissue sections were stained with the Eubacteria probe (white) **(A)** and custom-designed RNAscope probes against the 16S ribosomal RNA gene of bacteria belonging to the Bacillota Phylum (Magenta), Actinomycetota (cyan) **(B)** or to the *Streptococcus* genus (yellow) **(C)**, alongside immunohistochemistry against E-cadherin (red) and nuclear DAPI stain. Whole slide scans of stained patient tissue sections were obtained using a Vectra whole slide scanner (40x). Images were spectrally unmixed and viewed in Visiopharm.

## Discussion

Our data show that the modified Gram stain is superior in visualising high load of microbial biomass to the standard H&E stain (Figure 1-2). However, H&E is superior in identifying *H. pylori* in gastric tissue sections. This finding indicates that to visualise both *H. pylori* and non-*H. pylori* bacteria in the same clinical tissue sample, consecutive sections should be stained with H&E and the modified Gram stain to reveal *H. pylori* and non-*H. pylori* bacteria, respectively. Alternatively, the modified Gram stain should be further optimised to detect *H. pylori*. Nonetheless, given our recent study (5) in which we identified non-*H. pylori* bacteria invading the gastric lamina propria in *H. pylori-*associated gastric carcinogenesis, we have also shown here that the modified Gram Stain can be used to select samples with distinct bacterial morphologies, which validates the use of this tissue sample for downstream applications (Figure 3). An example of this could be laser capture microdissection (LCM) and sequencing to identify bacterial species in target microniches. However, LCM and sequencing of isolated bacterial DNA is a technically challenging and expensive approach (11, 12). Therefore, this initial screening for bacterial microniches with the modified Gram stain ensures success in resolving the bacterial identity within target microbial communities.

Here, given the recent interest of *S. anginosus* in the development of gastric cancer (10), we used a Phylum-specific probe against Bacillota (Firmicutes) and a *Streptococcus-*specific genus probe that detects most *Streptococci* species. Using consecutive sections of a gastric adenocarcinoma tissue, we were able to validate with both the modified Gram stain and custom-designed RNAScope probes towards Kingdom, Phyla and Genus probes that this tissue contained a diverse bacterial community, most likely as a multi-species biofilm (Figure 4A-C). However, further studies are required to fully interrogate the prevalence of these multi-species biofilms in gastric adenocarcinoma tissue in large patient cohorts and with three-dimensional reconstruction to ensure potential biofilm communities are not being missed by the staining of only single sections from large cancerous tissue. Nonetheless, this data validates the approach of combining RNAscope technologies with immunohistochemistry to visualise host-polymicrobial interactions in complex tissue microenvironments. Furthermore, given the interest in intratumoral microbiomes (13), this paper reports a range of histological tools to determine the validity of the intratumoral microbiome, which are independent of sequencing technologies.

In summary, we have presented both expensive/time-consuming and cheap/quick histopathological tools for the visualisation of the microbiota in tissue samples. This provides the opportunity for microbiome scientists to either view a ‘snapshot’ of bacterial communities before sequencing, or validate existing bioinformatic data of mucosal microbiomes by visualising top hits *in situ*. This has the potential to unlock new biology underpinning host-microbiota relationships, as we have previously shown in the context of gastric carcinogenesis (5).

## Acknowledgments

We thank the patients for providing tissue samples. We thank the Human Biomaterials Resource Centre for providing full ethical approval (#17-285). We thank the Royal Society (RGS\R2\192,312), the University of Birmingham Cancer Research UK Centre Development Fund (A.E.R-P.), Cancer Research UK (EDDPMA-Nov21\100,008) (A.E.R-P.) and the Medical Research Council IMPACT Doctoral Training Programme (H.J.G. and A.E.R-P.) for funding. We also thank Gareth Bicknell, Michael Russell, Emma Hurlestone, Shalom Simende and Shannon Smallman at HBRC for patient sample retrieval. We thank Dr. Gary Reynolds for tissue sample sectioning. We thank Joe Flint at Birmingham Tissue Analytics for whole slide scanning.

